# Chronic effects of short-term aerobic and anaerobic physical training on the ventral prostate of adult rats

**DOI:** 10.1101/2021.02.04.429709

**Authors:** Rodrigo Miranda Ramos Borges, Emerson Souza da Rocha, Edila Monteiro de Andrade, Nagaywer Edno da Silva Nazaré, Paulo Afonso Ortega Araújo, Pedro Nogarotto Cembraneli, Thalles Fernando Rocha Ruiz, Simone Jacovaci Colleta, Carla Patricia Carlos, Vanessa Belentani Marques, Sebastião Roberto Taboga, Fabiana de Campos Gomes, João Simão de Melo-Neto

## Abstract

**Aims:** To analyze the chronic effects of short-term aerobic and anaerobic physical training on prostate compartments, extracellular matrix, microvascularization, TGFβ, cyclooxygenase-2, inducible nitric oxide synthase (iNOS) and macrophage immunostaining, and ventral prostate histopathology in adult rats.

**Main methods:** Fifteen male rats (90 days old) were divided into three groups (n = 5/group): sedentary, aerobic (AE) (swimming), and anaerobic physical training (AN) (jumping), performed three days per week, for 8-weeks. The animals were sacrificed within 21 weeks of age. The ventral prostate was collected, weighed, and processed for histological and immunohistochemical analyses.

**Key findings:** Our results show that AE increases blood capillaries and reduces the percentage and increases the thickness of smooth muscle. AN promotes increased iNOS levels in the prostatic tissue, and both modalities reduce TGFβ and elastic fibers, in addition to being protective against benign prostatic hyperplasia and atrophy.

**Significance:** Different physical training modalities can activate specific mechanisms that modify the prostate environment.

## 1. Introduction

The prostate is an accessory sex gland that produces and stores a wide range of nutrients essential for sperm survival and viability [1]. In rats, it is organized and anatomically delimited surrounding the pelvic urethra and is divided into four pairs of lobes (anterior, ventral, lateral, and dorsal), forming the prostatic functional unit with histological characteristics and production of specific secretions [2, 3]. However, the ventral lobe has been studied more extensively because it is more responsive to hormonal modulation [4, 5].

The prostate is divided into three main compartments: the epithelium, lumen, and stroma. In the epithelium, the prostatic parenchyma produces secretions that are released in the lumens of the prostatic ducts. The stroma of the prostate consists of a complex arrangement of smooth muscle cells, extracellular matrix, nerve endings, fibroblasts, myofibroblasts, and blood vessels [6, 7]. The stromal components are closely associated with the prostatic epithelium; thus, the stroma acts in the regulation of epithelial proliferation and differentiation for the maintenance of tissue homeostasis. Microvascularization of the prostatic extracellular matrix undergoes constant remodeling, and scientific evidence shows that vascular changes trigger potential prostatic pathologies, reducing blood flow and decreasing the response to tissue repair [8]. Thus, an imbalance in the interaction between the stroma and the epithelium can lead to pathologies, with the development of a new microenvironment [9].

Androgens are one of the most important factors in homeostasis and the development of prostate tissue. Therefore, possible changes in hormonal levels can influence stromal remodeling, resulting in tissue adaptations in muscle cells and elastic and collagen fibers that may be present in pathological processes [10, 11].

Transforming growth factor-beta (TGFβ) is a polypeptide responsible for the control of differentiation, cell proliferation, development, and tissue repair [12]. The TGFβ Superfamily comprises 33 members, with the most common ones in mammals being TGFβ1, TGFβ2, and TGFβ3 [13]. The function of TGFβ in the prostate is related to prostate growth and angiogenesis, and it has been shown to be associated with prostate cancer [14]. It accumulates in stromal cells, contributing to the progression of proliferative lesions that occur in the prostate [15]. However, Lee et al. (1999) pointed out that the mode of action of TGFβ in prostatic tissue is variable [14].

COX-2 and iNOS are important inflammatory mediators involved in cancer-related inflammation [16]. Furthermore, at the tissue level, it is possible to evaluate the overexpression of enzymes associated with the progression of malignant prostatic pathologies, such as COX-2, which can be released by macrophages [17]. This enzyme is largely related to increased multiplication, migration, inflammation, angiogenesis, and metastasis of cancer cells related to inflammatory infiltrates and is a major component of several malignant carcinomas, including prostate cancer [18, 19]. COX-2-derived prostaglandins can increase tissue-resident macrophages [20]. Nitric oxide synthase (NOS) is a family of enzymes responsible for catalyzing nitric oxide, a potent vasodilator. iNOS is an enzyme of the NOS family present in inflammatory processes when there is a marked production of nitric oxide [21], and a positive correlation with macrophages has been demonstrated [22]. Under pathological conditions, iNOS is considered a harmful enzyme for tissues [21], associated with increased mortality in cases of prostate cancer with increased iNOS in the stroma [23].

Therapeutic possibilities for prostatic adenocarcinoma are limited, with androgenic suppression as the focus, aiming at the involution of pathologies [24, 25]. In this study, physical exercise was considered for use as a treatment option with significant results, as it affects the endocrine reproductive system in humans.

Treatment tools for prostatic adenocarcinoma are limited, highlighting the importance of androgen-deprivation therapy, which contributes to tumor regression [25]. The hypothalamic-pituitary-gonadal axis regulates the synthesis of androgens. The organs of the genital system, including the prostate, an androgen-dependent organ, are influenced by these hormones [26, 27]. This can be further complemented by physical exercise, which can reduce testosterone levels, depending on the intensity, duration, and regularity of the activity [26, 28].

When physical activity is performed in a structured, planned, and mainly progressive way, it is called physical training, which can be classified according to its objective. The effects of exercise depend on this training and are controlled by the amount, frequency, duration, and intensity of training [29].

Resistance training is performed with a complement of load, with weight, body mass, or elastic devices [30]. During this exercise, the muscle must exert force against resistance, with an effort above the aerobic capacity of the muscle, with contractions that lead to fatigue. Therefore, this exercise is of high intensity and short duration [31] in most cases. This type of exercise is considered to be anaerobic [32], because energy generation depends mainly on anaerobic energy pathways, unlike aerobic exercise, which aims to adapt body systems to maintain the oxygen demand for a tissue during exercise training. According to the American College of Sports Medicine, any activity that uses large muscle groups, can be maintained continuously, and is rhythmic can be considered as aerobic exercise [31].

As reviewed by Friedenreich, Neilson, and Lynch (2010) [33], physical activity reduces the risk of development of prostate cancer, and intense exercise can modify hormone levels. Resistance exercise has also been reported to decrease circulating testosterone levels [26]. However, another study [34] pointed out an increase in circulating testosterone and DHT levels in animals, demonstrating discrepancies in the literature.

A systematic review [35] to verify the effects of aerobic exercise on the incidence, progression, and metastasis of prostate cancer found significant heterogeneity in the study methodologies, making it difficult to determine the relationship between these variables. There is little research [36] on experimental models involving aerobic and anaerobic physical training. In studies using mouse models of prostate cancer, it was observed that this type of exercise can modulate the tumor microenvironment, reducing hypoxia, leading to less phenotypic aggression, which can favor a better prognosis for the patient [36]. However, the mechanisms involved in the prevention or appearance of pathological changes need to be elucidated.

Therefore, this study aimed to analyze the chronic effects of short-term aerobic and anaerobic physical training on prostate compartments, extracellular matrix, microvascularization, TGFβ, COX-2, iNOS and macrophage immunostaining, and histopathology of the ventral prostate of adult rats.

## 2. Methods

### 2.1. Animals and experimental environment

Fifteen male Sprague-Dawley rats (90 days old) were obtained from the Multidisciplinary Center for Biological Research of the University of Campinas (CEMIB/UNICAMP). Sprague-Dawley rats are especially used in biological research related to oncology because they are more susceptible to pathological changes [37] and are considered an appropriate experimental model for this study. The age of the animals was determined based on previous studies [38-40].

Rats were housed in the Laboratory of Experimental Research of Faceres School of Medicine (FACERES) under adequate lighting (12-hour light-dark cycle) and temperature conditions (22 ± 2°C). Throughout the experimental period, the animals had free access to a common solid diet (Labina Purina®) and water. All rats were weighed weekly, and food and water intake was measured daily. All experiments and surgical procedures in this study were approved by the Ethics Committee for Animal Use of the FACERES (protocol number 02/2016).

### 2.2. Experimental groups

Animals were randomly distributed into three experimental groups (five rats each): sedentary (SD), anaerobic (AN), or aerobic physical training (AE). The software GPower (3.1, Franz Faul, Universitat Kiel, Germany, 2007) was used to calculate the statistical power of the sample. Elastic fiber data (%) were randomly selected to calculate the effect size f (0.742) of the F test analyses. The statistical power obtained for the F tests was 0.614, considering an α error probability of 0.05. To calculate the statistical power of the chi-square test (χ^2^), the noncentrality parameter (λ) = 11,935 was initially calculated. It was assumed as the basis: i) Generical χ^2^; ii) α probability error = 0.05; iii) Df = 4. The statistical power obtained for the χ^2^ test was 0.799.

### 2.3. Experimental protocol

The experimental protocol was carried out 3 days per week, for 8 weeks in the early evening (18h 00), after the beginning of the dark cycle. The 8-week period was chosen due to the duration of the spermatogenesis cycle (48–56 days) [41, 42], which is the minimum period for changes to occur in the hypothalamus-pituitary-gonad axis. To simulate stress in a liquid environment, the animals in the SD groups were placed in a container containing shallow water (±30 °C). The animals were subjected to resistance physical training (anaerobic) and swimming (aerobic).

#### 2.3.1. Resistance physical training (anaerobic)

The AN group was subjected to jumping sessions in a cylindrical PVC containing water at a temperature of ± 30 °C and a depth of 38 cm [43]. Initially, the animals were subjected to a period of weight overload adaptation, following which the animals were subjected to a protocol adapted from Cunha et al. (2005) [44] and de Melo Neto et al. (2015) [39], with weight overload placed in a vest attached to the ventral region of the thorax (Table 1). The material used as an overload was a screw and washer of iron. The vest material was cotton fabric with Velcro. This anaerobic physical training protocol was previously validated by evaluating the blood lactate concentration in a study performed by Tanno et al. (2006) [45].

**Table 1.**
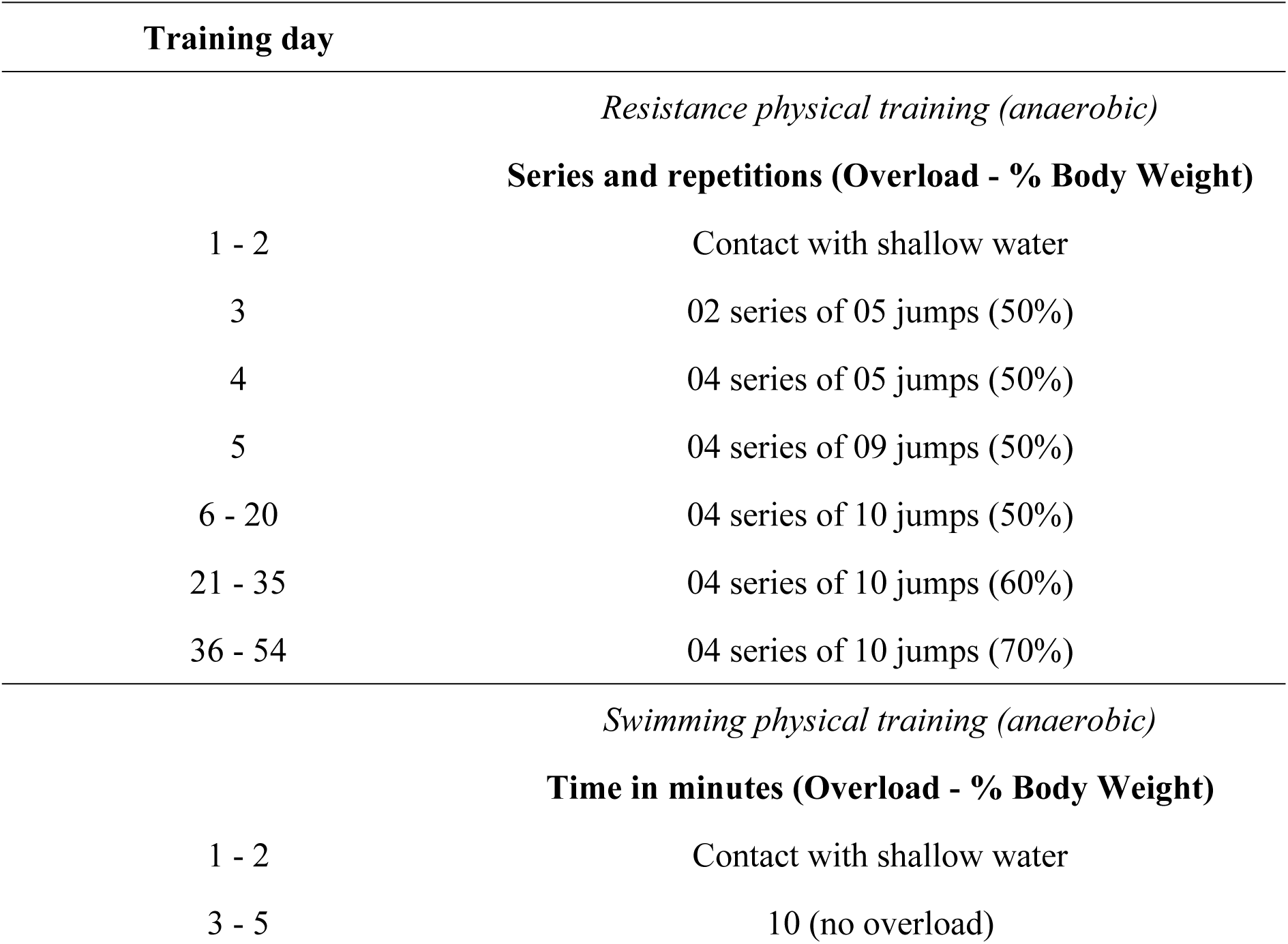
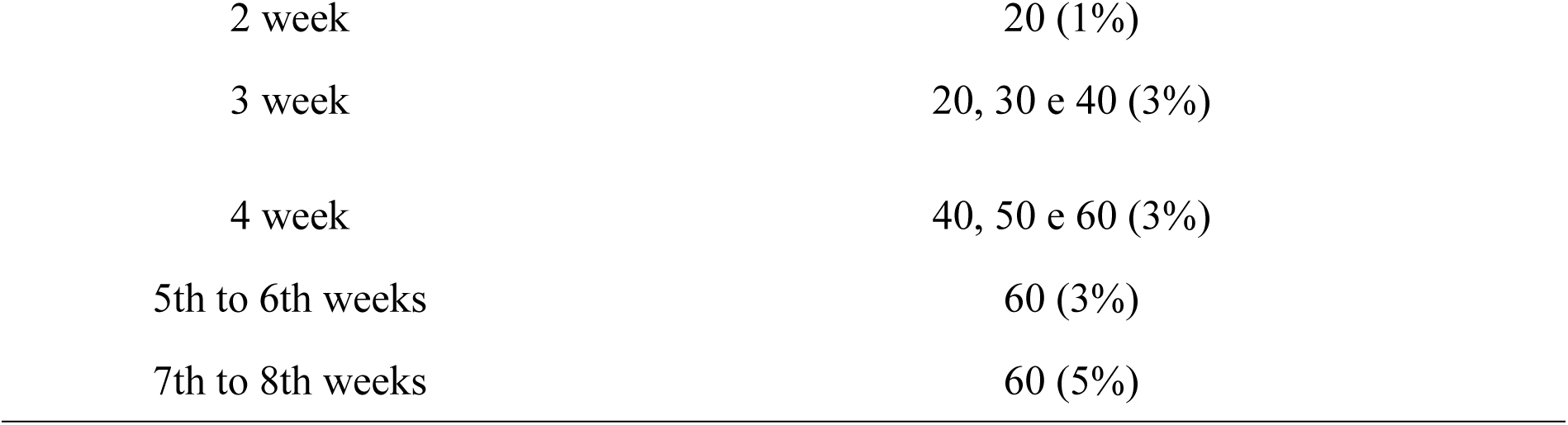
The protocol of anaerobic and aerobic physical training.

| Training day |  |
| --- | --- |
| *Resistance physical training (anaerobic)* |  |
| **Series and repetitions (Overload - % Body Weight)** |  |
| 1 - 2 | Contact with shallow water |
| 3 | 02 series of 05 jumps (50%) |
| 4 | 04 series of 05 jumps (50%) |
| 5 | 04 series of 09 jumps (50%) |
| 6 - 20 | 04 series of 10 jumps (50%) |
| 21 - 35 | 04 series of 10 jumps (60%) |
| 36 - 54 | 04 series of 10 jumps (70%) |
| *Swimming physical training (anaerobic)* |  |
| **Time in minutes (Overload - % Body Weight)** |  |
| 1 - 2 | Contact with shallow water |
| 3 - 5 | 10 (no overload) |
| 2 week | 20 (1%) |
| 3 week | 20, 30 e 40 (3%) |
| 4 week | 40, 50 e 60 (3%) |
| 5th to 6th weeks | 60 (3%) |
| 7th to 8th weeks | 60 (5%) |

#### 2.3.2. Swimming physical training (aerobic)

The animals were subjected to swimming physical training for over 8 weeks [46], three times/week [47]. The AE group was subjected to progressive aerobic training in a water tank (110 × 70 × 22 cm), filled with 50 cm of water at a temperature of 30±2 °C. The animals were previously subjected to a period of adaptation to water. Adaptation took place during the first five days. During the protocol period, the animals swam using washers attached to an elastic band on their tails to generate overload. The training sessions were progressively organized according to Table 1, as proposed in previous studies [39, 48].

### 2.4. Euthanasia, tissue collection and processing

Animals were euthanized within 21 weeks of age, 48 h after the end of the experiment, through a lethal dose (100 mg/kg) of sodium thiopental (Tiopental®) administered by intraperitoneal injection. The prostate tissue was collected and weighed. Tissues were fixed in 4% paraformaldehyde in phosphate-buffered saline (PBS) pH 7.4, processed, embedded in paraffin, cut into 3-μm thick sections, and stained. Morphometric, stereological, histopathological, and immunohistochemical analyses were performed.

### 2.5 Morphometric and stereological analyses

For morphometric and stereological analysis, five sections were used per group, with 40 fields per group selected at random (objective magnification 40x) being analyzed. Histological sections stained with hematoxylin and eosin (H&E) were used to analyze the prostate compartments (epithelium, lumen, muscle, and non-muscular stroma), epithelium height (μm), and smooth muscle thickness (μm), these being analyzed in 40 fields/group selected at random. Gomori reticulin staining was used to investigate reticular fibers (type III collagen) and Verhoeff for elastic fibers to analyze cellular matrices.

The relative volumes (%) of the extracellular matrices and the different compartments were evaluated using stereological analysis. For this, we followed the method proposed by Weibel et al. (1966) [49], using a 418-point grid in the fields.

### 2.6 Microvascularization analysis

For each animal, 40 fields randomly selected per group (eight per animal) were examined under a microscope (40× magnification), counting the number of capillaries and arterial and venous blood vessels.

### 2.7 Immunohistochemistry analyses of *TGFβ, COX-2, iNOS, and ED1*

Deparaffinized cuts were subjected to the recovery of antigenic activity in sodium citrate buffer in the microwave, and endogenous tissue peroxidase block and pre-primary protein block were performed. The primary antibodies used for incubation were anti-TGFβ 1/2/3 [H-112] (dilution 1: 80, Santa Cruz Biotechnology, Inc., sc-7892), anti-COX2 antibody [EP1978Y] (dilution 1: 500, Abcam), NOS2 (C-11): (dilution 1: 50, Santa Cruz Biotechnology, Inc.inc sc-7271), and anti-ED1 [MCA341R] (dilution 1: 500, Bio-Rad Laboratories, Inc). After overnight incubation with primary antibodies, the sections were washed in PBS and incubated with biotinylated secondary antibodies and HRP-streptavidin, using the Histostain SP Broad Spectrum kit (Invitrogen Cat # 959943 B) for 1 h at 37 °C. The slides were then treated with diaminobenzidine (DAB) (ref. 002014, Invitrogen), contrasted with hematoxylin, assembled with coverslips, observed, and analyzed. A negative control was performed for all immune reactions to verify specificity.

Twelve fields within the tissue area were randomly assigned to each group to evaluate the percentage (%) of immunoreactivity. For analysis of TGFβ, COX-2, and iNOS, the fields were analyzed using the Image J 1.47 Windows version software (National Institutes of Health, USA), according to Ruifrok and Johnston’s method (2001) [50]. During immunohistochemistry analysis, a fixed threshold was set to obtain the percentage of immunostained tissue area corresponding to the analyzed proteins [51]. Quantification of the ED-1 reaction was performed by counting the number of positive cells by field.

### 2.8. Photo capture

Analyses were performed on images acquired using a Zeiss Primo Star microscope coupled to a Zeiss Axiocam 105 color camera (objective magnification 40 ×) using the Zen Lite 2.3 software (Zeiss).

### 2.9 Histopathological analysis

Histopathological analysis was performed on H&E-stained slides to evaluate the presence of histopathological lesions, as described by Creasy et al. [52]. Tissue sections were analyzed using an objective magnification of 10×, 20×, and 40×.

### 10. Statistical analysis

Data were analyzed using descriptive and inferential statistics. Descriptive results are expressed according to normal distribution. Parametric data are expressed as mean ± standard deviation, while non-parametric data are described as median, minimum, maximum, and quartiles (Q1, Q3). The Shapiro-Wilk test was used to analyze the normality of the data. During intergroup comparisons, the following tests were applied: analysis of variance (ANOVA) with post-hoc Tukey’s test (parametric data), Kruskal-Wallis test, and post-hoc Dunn’s test (non-parametric data). For the presence of pathological processes, the association was analyzed using the chi-square test (χ^2^). Statistical significance was set at p ≤ 0.05. Statistical analysis was performed using the Instat software (version 3.0, GraphPad, Inc., San Diego, CA, USA).

## 3. Results

### 3.1 Biometric parameters

Table 2 shows the animals’ body weight, absolute and relative weight of the ventral prostate, and feed consumption and water intake by the animals during the experimental period. Anaerobic physical training promoted a reduction in feed consumption and water intake, final body weight, and body weight variation.

**Table 2.** Body weight, absolute and relative weight of the ventral prostate, consumption of feed, and intake of water in the different experimental groups.

|  | SD | AN | AE | F or H | *p* * |
| --- | --- | --- | --- | --- | --- |
| **Feed consumption (g)** | 1400 ± 51.95<sup>a</sup> | 1289.4 ± 10.06<sup>b</sup> | 1412.8 ± 86.10<sup>a</sup> | 6.738 | 0.0109* |
| **Water intake (ml)** | 2439 ± 255.4 | 2175 ± 67.89<sup>a</sup> | 2624 ± 174.4<sup>b</sup> | 7.621 | 0.0073* |
| **Initial body weight (g)** | 495 ± 36.40 | 511.3 ± 11.39 | 514 ± 54.47 | 0.3578 | 0.7064 |
| **Final body weight (g)** | 525<br>(136.50;199.56)<sup>a</sup> | 481 (87.80;<br>155.09)<sup>b</sup> | 515 (111.71;<br>210.30) | 7.645 | 0.0138* |
| **Body weight variation (final - initial)** | 33 ± 21.68<sup>a</sup> | -27.5 ± 33.45<sup>b</sup> | 13 ± 10.37<sup>a</sup> | 8.402 | 0.0052* |
| **Absolute weight of the ventral prostate (g)** | 0.395 ± 0.186 | 0.352 ± 0.037 | 0.371 ± 0.106 | 0.1450 | 0.8665 |
| **Relative weight of the ventral prostate (absolute ventral prostate weight / final body weight)** | 0.074 ± 0.034 | 0.071 ± 0.007 | 0.071 ± 0.018 | 0.0365 | 0.9643 |

\**p* < 0.05, one-way ANOVA + Tukey's post-test ("a" compared to "b") or Kruskal-Wallis + Dunn's post-test. SD: sedentary; AN: anaerobic; AE: aerobic.

### 3.2. Compartments, morphometry, and micro vascularization

Data related to compartments, morphometric, and microvascular analyses of the ventral prostate are shown in Table 3 and Fig. 1. Aerobic physical training reduced the percentage of tissue smooth muscle but increased its thickness. Aerobic physical training reduced the number of arterial and venous blood vessels and promoted an increase in blood capillaries within the microvasculature.

**Table 3.** Compartments, morphometry, and microvascularization in the different experimental groups

|  | **SD** | **AN** | **AE** | **F or H** | ***p*\*** |
| --- | --- | --- | --- | --- | --- |
| *Compartments* |  |  |  |  |  |
| **Epithelium (%)** | 56 (42;79) | 62.5 (52;78) | 57.5 (48;76) | 0.076 | 0.9627 |
| **Lumen (%)** | 217 (161;262) | 217.5 (176;233) | 244.5 (208;272) | 1.842 | 0.3981 |
| **Non-muscular stroma (%)** | 101.5 (62;144) | 112.5 (83;181) | 120 (45;151) | 0.965 | 0.6171 |
| **Stroma (smooth muscle) (%)** | 20 (17;24)<sup>a</sup> | 21 (20;24)<sup>a</sup> | 14.5 (12;17)<sup>b</sup> | 12.75 | 0.0017* |
| *Morphometry* |  |  |  |  |  |
| **Epithelium height (µm)** | 104.1 (95.67;113.6) | 114 (106.1;124.2) | 108.7 (99.73;122.40) | 0.7001 | 0.7047 |
| **Thickness of the layer of smooth muscle (µm)** | 24.19 (21.63;27.00)<sup>a</sup> | 26.83 (24.19;29.55) | 27.66 (25.63;30.59)<sup>b</sup> | 6.982 | 0.0305* |
| *Microvasculature* |  |  |  |  |  |
| **Capillaries (n / field)** | 1.5 (1;2)<sup>a</sup> | 2.5 (1;3) | 3 (2;4)<sup>b</sup> | 10.92 | 0.0043* |
| **Arterial blood vessels (n / field)** | 0.00 (0;1)<sup>a</sup> | 0.00 (0;0) | 0.00 (0;0)<sup>b</sup> | 6.669 | 0.0356* |
| **Venus blood vessels (n / field)** | 1.00 (0;1)<sup>a</sup> | 0.00 (0;1) | 0.00 (0;0)<sup>b</sup> | 12.39 | 0.002* |
| *Extracellular matrix* |  |  |  |  |  |
| **Reticular fibers (%)** | 27.5 (25;31) | 24.5 (18;29) | 24.5 (21;28) | 5.699 | 0.0579 |
| **Elastic fibers (%)** | 23.95 ± 8.33<sup>a</sup> | 14.95 ± 5.00<sup>b</sup> | 14.33 ± 4.44<sup>b</sup> | 30.51 | <0.0001* |

\*p < 0.05, one-way ANOVA + Tukey's post-test ("a" compared to "b") or Kruskal-Wallis + Dunn's post-test ("a" compared to "b"). SD: sedentary; AN: anaerobic; AE: aerobic.

**Fig. 1.**
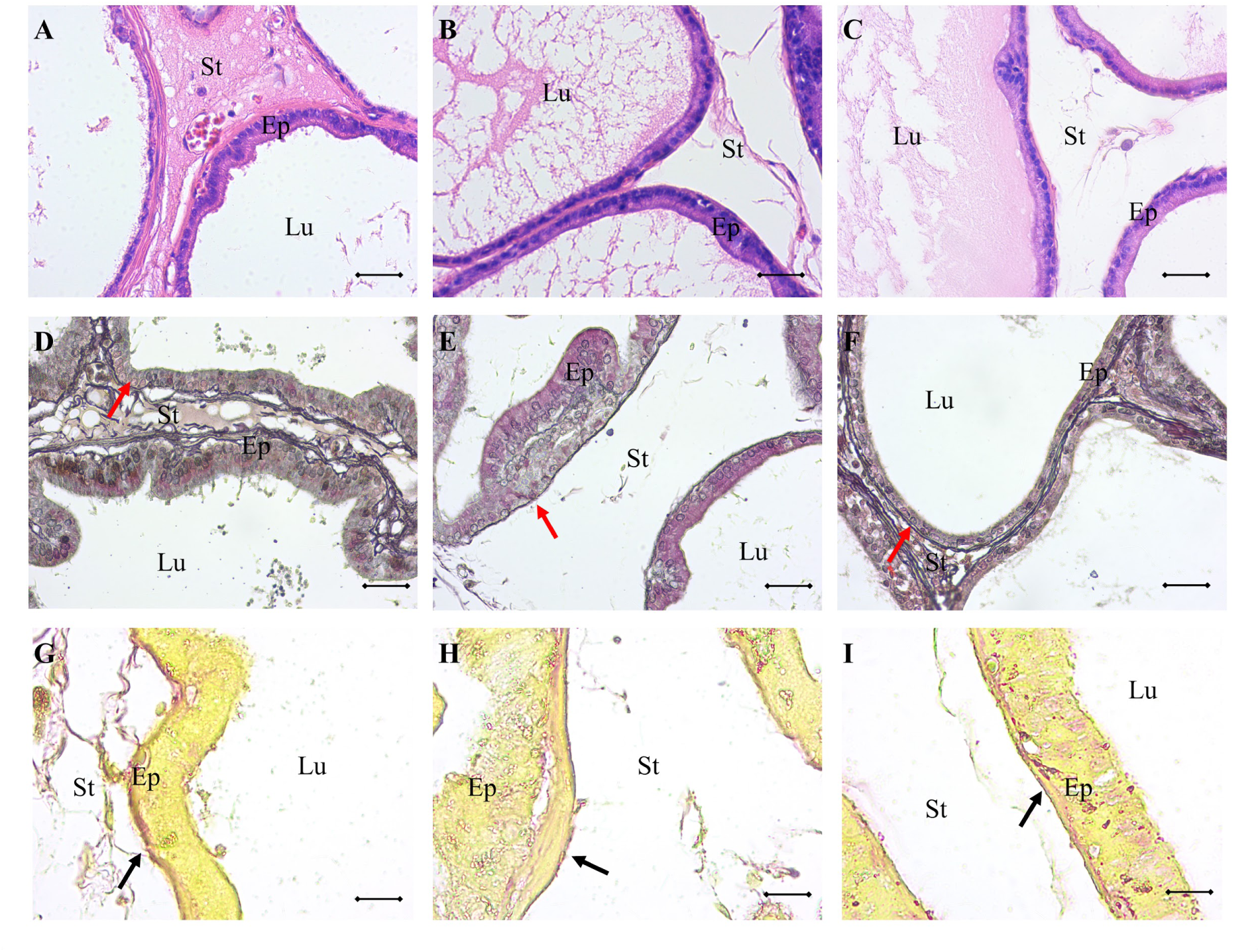
Normal histology (A–C), reticular fibers (D–F) and elastic fibers (G–I) in Groups SD (sedentary) (A, D, G), AN (anaerobic) (B, E, H) and AE (aerobic) (C, F, I). Ep: epithelium; St: stroma; Lu: Lumen. Scale bar: 50 μm.

**Fig. 2.**
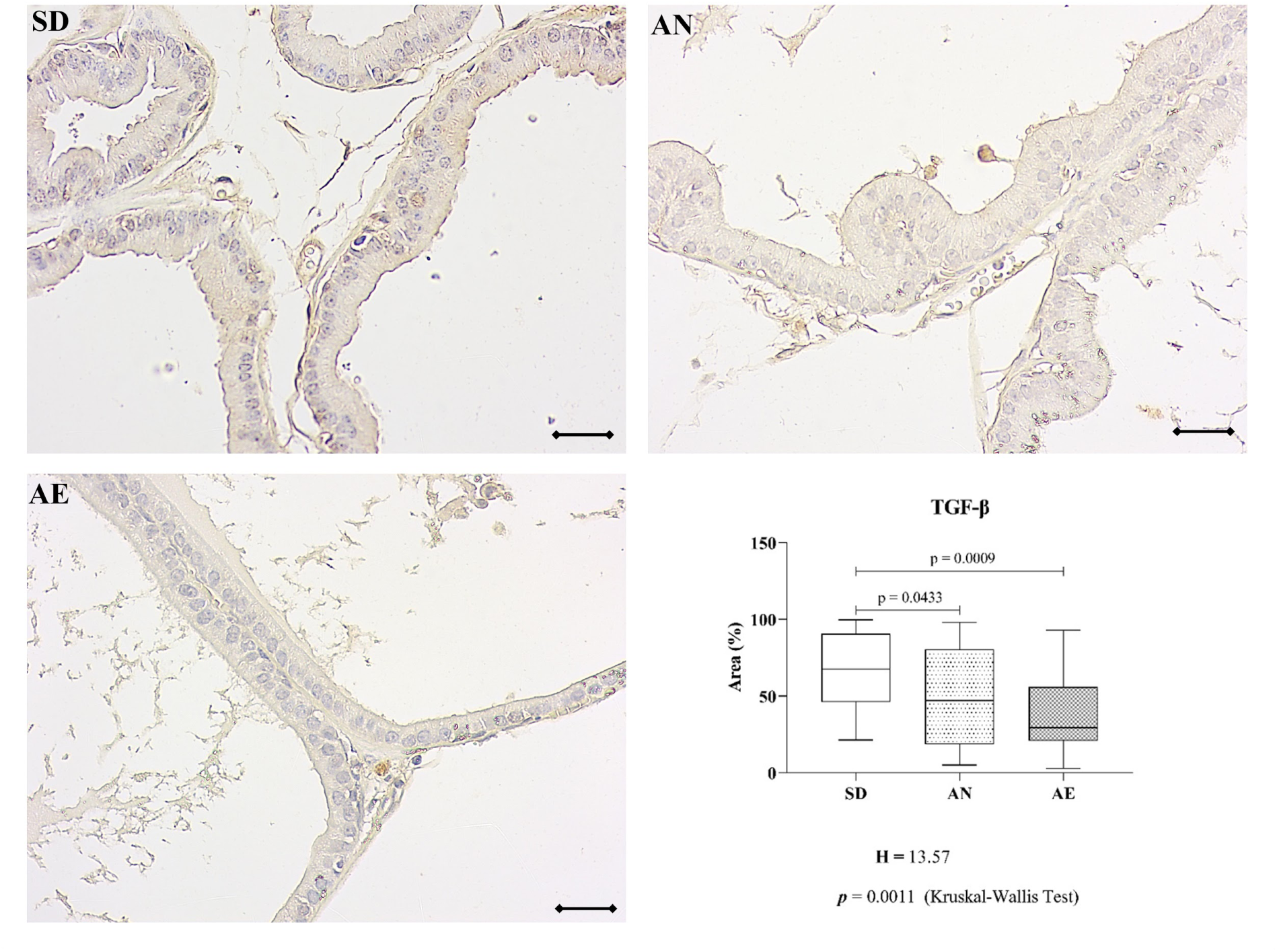
Immunostaining of transforming growth factor-beta (TGFβ) in groups: sedentary (SD), anaerobic (AN), and aerobic (AE) and percentage of the immunoreactive area (Graphic). Scale bar: 50 μm.

### 3.3. Extracellular matrix

Data related to extracellular matrix analyses of the ventral prostate are shown in Table 3 and Fig. 1. Both anaerobic and aerobic physical training reduced elastic fibers. There were no differences in reticular fibers after anaerobic and aerobic physical training.

### 3.4. Immunoreactive area of TGFβ, COX-2, iNOS, and ED1

Fig. 3, 4, 5, and 6 show the immunostaining of TGFβ, COX-2, iNOS, and ED1. Physical training, regardless of modality, reduced TGFβ levels. Anaerobic treatment increased the iNOS levels. COX-2 and ED1 levels did not differ between the groups.

**Fig. 3.**
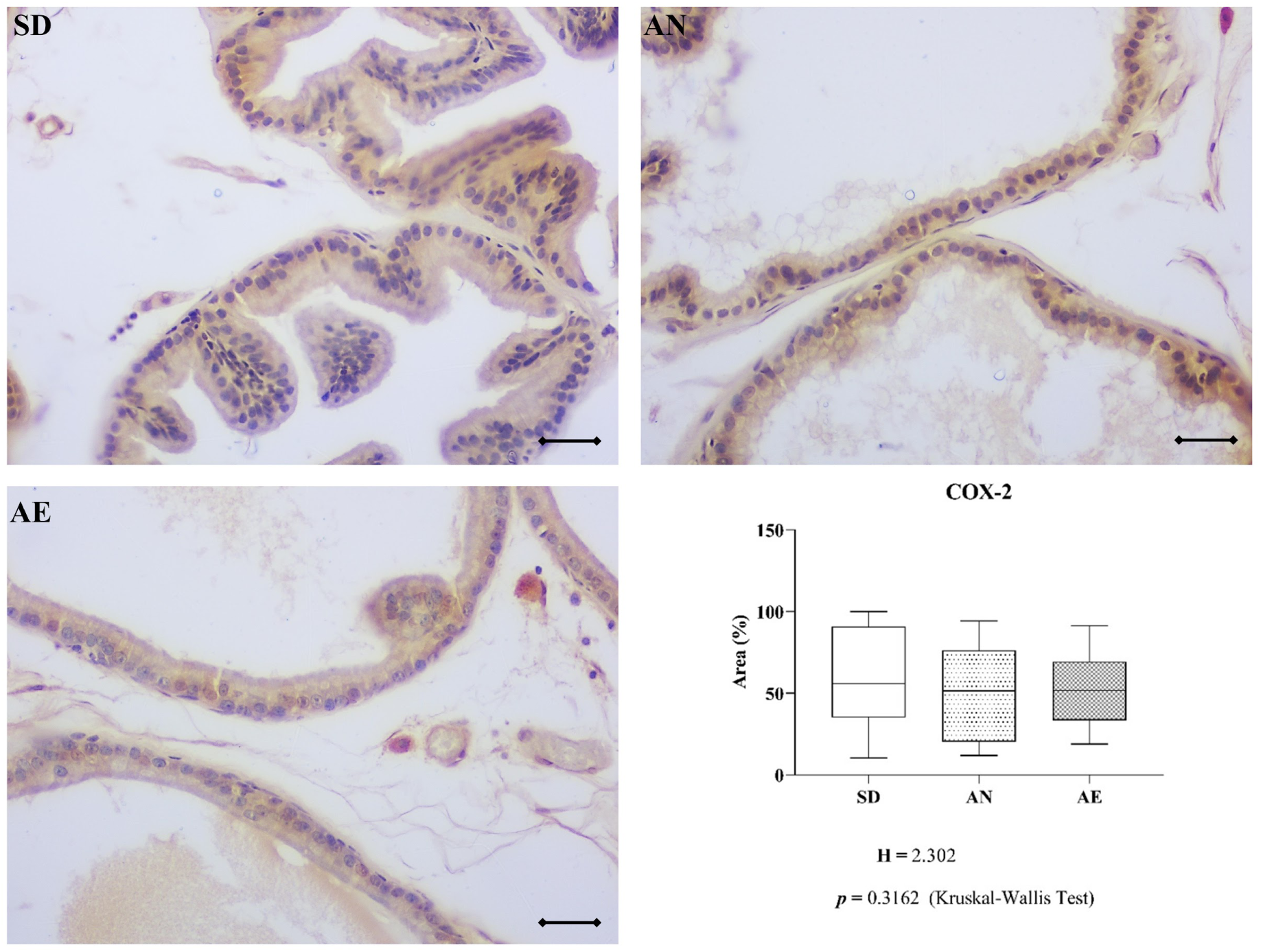
Immunostaining of cyclooxygenase-2 (COX-2) in groups: sedentary (SD), anaerobic (AN), and aerobic (AE) and percentage of the immunoreactive area (Graphic). Scale bar: 50 μm.

**Fig. 4.**
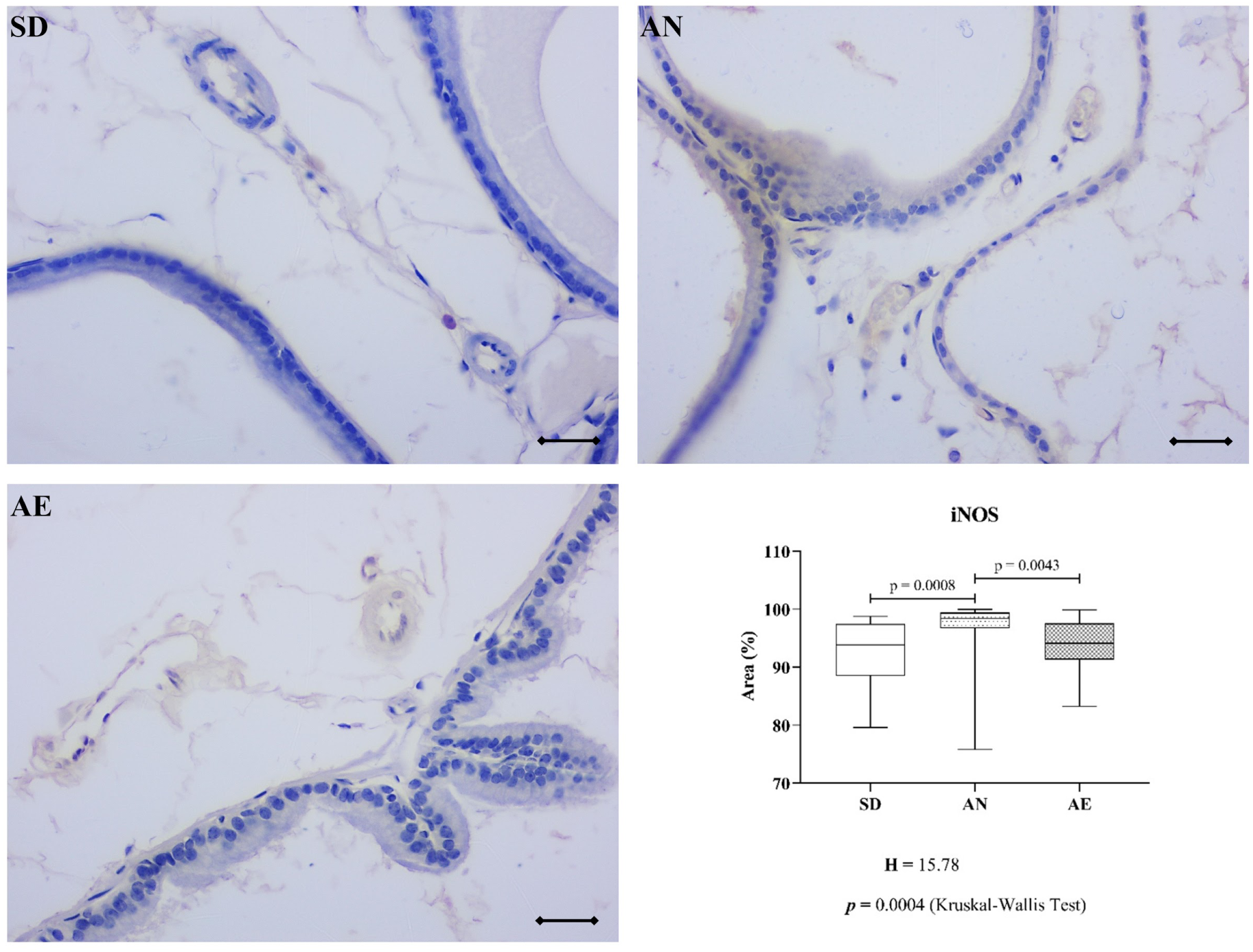
Immunostaining of induced nitric oxide synthase (iNOS) in groups: sedentary (SD), anaerobic (AN), and aerobic (AE) and percentage of the immunoreactive area (Graphic). Scale bar: 50 μm.

**Fig. 5.**
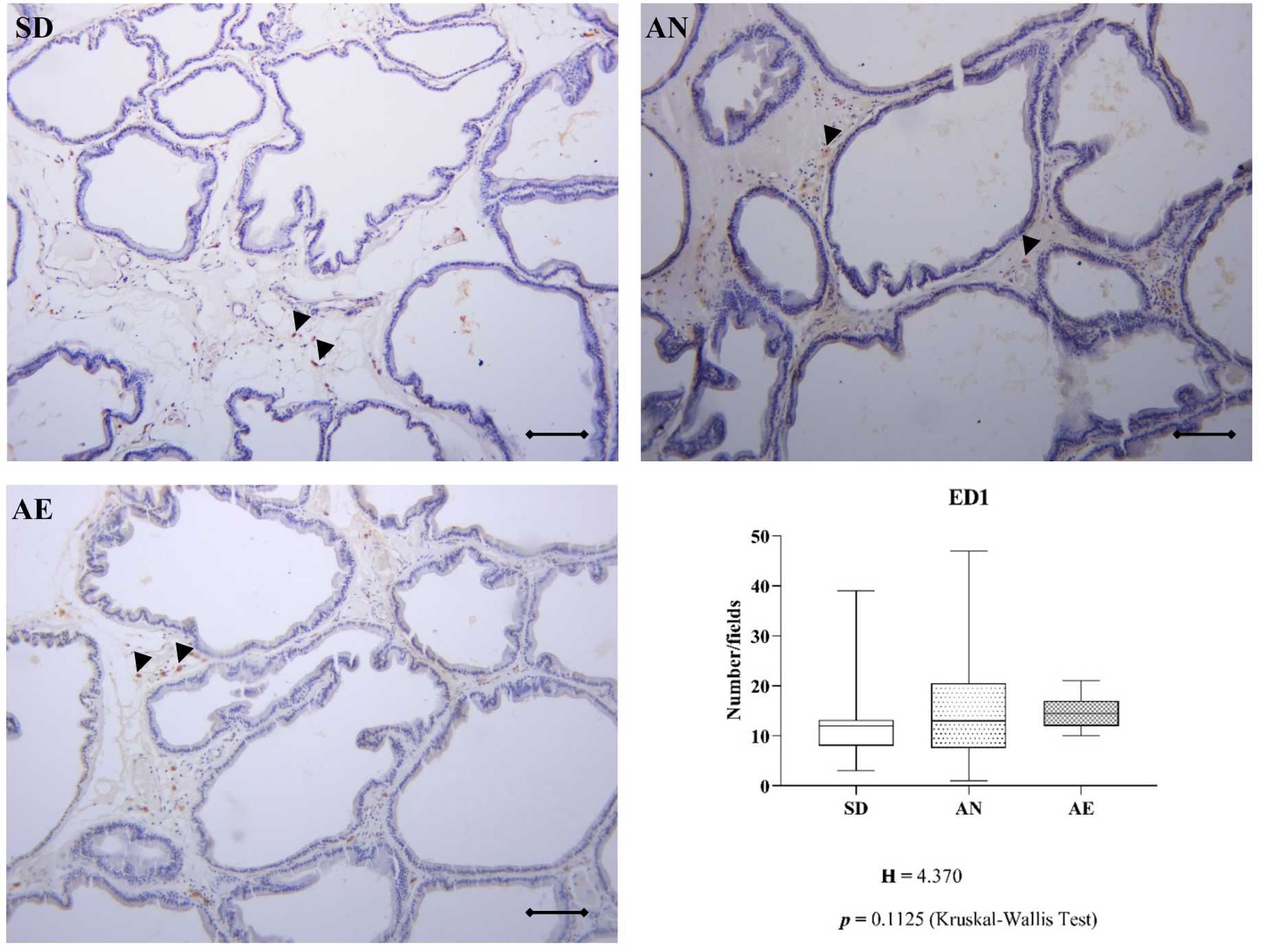
Immunostaining of macrophages (ED1) in groups: sedentary (SD), anaerobic (AN), and aerobic (AE) and percentage of the immunoreactive area (Graphic). Arrowhead: macrophages. Scale bar: 150 μm.

### 3.5. Histopathology

The histopathological analysis is shown in Table 4. Atrophy and benign prostatic hyperplasia were the only statistically significant factors, with protection in the AN group and AE.

**Table 4.**
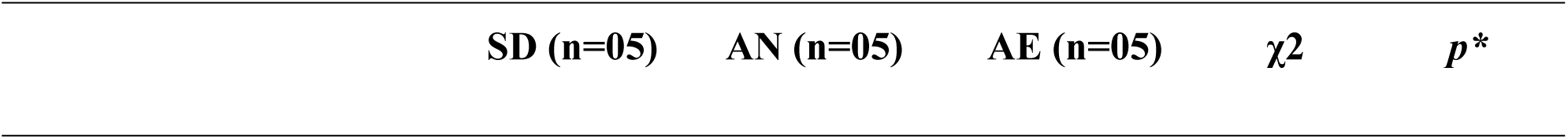
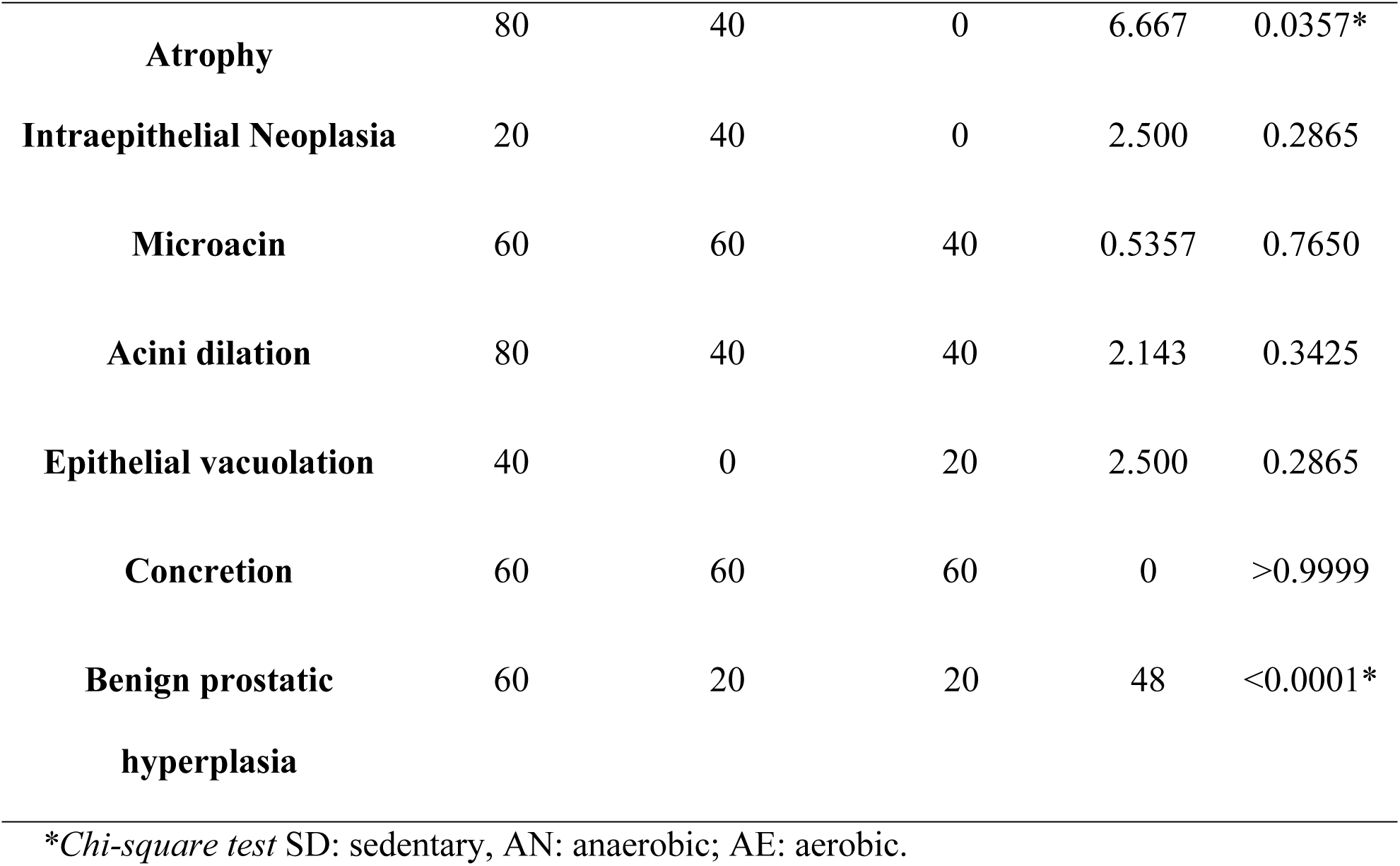
Histopathological analysis.

## 4. Discussion

Different types of physical exercise was able to influence the modulation of hormonal levels in different ways [26, 28]. Considering that the prostate is an androgen-dependent organ [26, 27], understanding the effects of exercise on this gland is extremely important. In these circumstances, we found that the animals that fulfilled the different exercise modalities had different effects from those that were not involved in this practice. The AE group presented adaptations in vascularization and smooth muscular tissue, while the AN group had reduced body weight, food consumption, water intake, and increased tissue iNOS levels. However, regardless of the modality, the practice of physical training resulted in the reduction of atrophy, benign prostatic hyperplasia, tissue TGFβ levels, and percentage of elastic fibers.

Pathological processes lead to increased cytokine levels in prostate tissue. TGFβ is an essential cytokine in reactive stroma activation processes, high levels of which can cause structural and functional changes in the extracellular matrix (ECM), and can also act on other types of cells such as epithelial, endothelial, and defense [53]. The decrease observed in our study can be a protective factor, as it negatively regulates the activation by pathological processes. In addition, we observed a reduction in the percentage of elastic tissue fibers. These fibers are influenced by TGFβ levels [54]. Simultaneous reduction of these tissue components after physical training has been demonstrated in the literature [55]. Changes in the ECM components contribute to the development of pathologies [56]. As a result, training has properties capable of generating positive tissue adaptations in prostate tissue.

Benign prostatic hyperplasia (BPH) has a strong influence on the pathogenesis of androgens, being the action induced by growth factors such as TGFβ, which is one of the main factors affecting proliferation, growth interruption, differentiation, and apoptosis of prostatic stromal cells. These diverse biological effects are directly correlated with the development of BPH, according to Lee et al. [57]. Thus, with the reduction of TGFβ levels by physical training, we can consider it a strong protective factor for this tissue adaptation observed in our study.

In addition, the atrophic glandular epithelium in the prostate was reduced in the animals that underwent physical training. Glandular atrophy is one of the main regenerative lesions of the prostate epithelium [58]. Exercise appears to be a regulatory mechanism for tissue dysfunction. The normalization of this alteration occurs through an increase in proliferative activity and maintenance of apoptotic activity levels [59, 60, 61].

The AN group showed a reduction in body weight, accompanied by decreased feed consumption and water intake. Intense training such as the anaerobic modality, along with high metabolic consumption, is associated with an increase in lipolysis and gluconeogenesis, depreciating fat reserves and glycogen. With this mechanism, a reduction in body weight occurs naturally if there is not enough caloric intake to compensate for the mobilization of reserves and consumption [62]. Changes in the eating habits of animals in the AN modality of our study suggested that the reduction in body weight was a consequence of decreased feed consumption and caloric intake [62].

The AN group showed a reduction in body weight, accompanied by a reduction in feed and water intake compared to sedentary rats. Intense anaerobic training is also responsible for increase in lipolysis and gluconeogenesis, mainly due to the reduction in food and calorie consumption, depriving more reserves of fat and glycogen [62], Conversely, aerobic physical training, due to its low intensity and high duration, is known to promote greater energy expenditure during exercise and for the strong oxidative component of fat and weight loss [31, 62]. Our study could not validate these results, as there was no statistical difference between the anaerobic and aerobic groups. The basic mechanism is that there is a reduction in body weight in case of insufficient calorie consumption to compensate for the mobilization and usage of reserves [62]. Observing the eating habits of animals in the AN modality of the study, it can be considered that these habits resulted in weight loss, as there was a reduction in feed consumption and caloric intake, consequently causing a reduction in body weight [62]. However, an alternative hypothesis is survival stress, which would have been perhaps more intense than physical stress itself, especially in the AN group [63].

In addition, the animals in the AN group also showed higher tissue iNOS levels. The relationship between iNOS inhibition and changes in body weight has been previously reported [64], corroborating our results. iNOS is not normally expressed and can be induced by tissue damage. Studies have shown that physical training can cause an increase in iNOS [65], supporting our findings.

After aerobic physical training, we observed a reduction in the percentage of muscle stroma and arterial and venous vascularization. Smooth muscle can influence vascularization and plays a crucial role in the mechanics of neovascularization. When the smooth muscle contracts, it generates folds in the vascular endothelium, stimulating its formation and supporting its patterns. Some studies on the development of vascularization have shown that a reduction in smooth muscle can cause a decrease in vascularization or even the development of tortuous and structurally non-functional vessels. Thus, the smooth muscle acts on the structure and formation of new vessels [66]. However, the decrease in smooth muscle was accompanied by an increase in its thickness, suggesting a compensatory mechanism.

In contrast, in our study, we observed an increase in the number of capillaries in the AE group. Aerobic physical training can cause adaptations in the structure and number of capillaries to accommodate higher aerobic demands and perfusion levels [67]. Additionally, it has great value as a non-pharmacological treatment of pathologies such as arterial hypertension, through its ability to regulate peripheral resistance and increased vascular tone [68]. This type of capillary angiogenesis has been suggested as a mechanism to improve the oxygen extraction capacity of tissues by increasing the area of diffusion [69, 70].

## 5. Conclusion

Both modalities are related to the reduction of TGFβ levels and elastic fibers, in addition to being protective factors for atrophy (mostly aerobic modality) and BPH. Only aerobic physical training for a short period of time reduces the percentage of smooth muscle and increases its thickness, modifies blood vascularization, and causes an increase in blood capillaries. In turn, the anaerobic modality increases iNOS levels in prostate tissue. Thus, this study can contribute to new therapeutic perspectives by understanding the mechanisms related to the effects of short-term physical training on the ventral prostate of adult rats.

## Acknowledgment

We are grateful to the Microscopy and Microanalysis Laboratory at Universidade Estadual Paulista / UNESP and the Clinical and Experimental Research Unit of the Urogenital System.

## Declaration of competing interest

The authors declare that there are no conflicts of interest

